# Membrane-binding of the bacterial dynamin-like protein DynA is essential for effective phage defence

**DOI:** 10.1101/2025.03.27.645684

**Authors:** Samia Shafqat, Urska Repnik, Marc Bramkamp

## Abstract

Bacterial dynamin-like proteins are large GTPases that play crucial roles in membrane dynamics. DynA in *B. subtilis*, a two-headed bacterial dynamin-like protein, possesses membrane-binding and tethering functions in trans. The formation of large DynA clusters on membranes in response to pore-forming antibiotics and phages demonstrates its potential role in maintaining membrane integrity under various environmental stresses. In this study, we identified the membrane-binding site of *B. subtilis* DynA within the D1 domain of the protein that includes positively charged lysine residues K360 and K367, as well as hydrophobic phenylalanine residues F363, F364, and F365. For experimental validation, recombinant proteins with mutations in the lysine or phenylalanine residues were produced and used in liposome binding assays. Non-conservative mutations lead to a complete loss of DynA’s membrane-binding capability. In vivo data showed that the membrane-binding mutants exhibit significantly increased susceptibility to phage infection compared to wild-type strains, further emphasizing the importance of DynA’s membrane interaction in conferring phage resistance. Our findings bridge the gap between the structural characteristics of DynA and its functional implications in maintaining membrane integrity and mediating phage resistance.

## Introduction

The dynamin superfamily of proteins is a large group that plays crucial roles in several essential cellular processes [1], including fission and fusion of biological membranes [2, 3], endocytosis and exocytosis [4], membrane remodeling [5], and immune responses to pathogens [6]. This large family is further divided into classical dynamins and dynamin-like proteins (DLPs), based mainly on sequence homology and notable structural features, including the highly conserved N-terminal GTPase domain, a central hinge domain, and an extended alpha-helical stalk domain [7]. The proteins are mechanochemical GTPases, which utilize the energy of GTP hydrolysis to remodel cellular membranes. They are over 60 kDa and can self-assemble on biological membranes, which activates their GTPase activity [8]. GTP binding and hydrolysis are mediated by four well-conserved motifs within the GTPase domain: G1, G2, G3, and G4 [1]. The G1 motif primarily binds to phosphates, the G2 motif, with a conserved threonine, interacts with Mg²⁺ ions to carry out GTP hydrolysis, the G3 motif also interacts with Mg²⁺ ions through aspartate, and the conserved glycine interacts with the γ-phosphate and the G4 motif interacts with the nucleotide base [9].

After the discovery of the first dynamin from bovine tissue and the shibire (ts) mutant in fly it became quickly clear that dynamin-like proteins are ubiquitous in eukaryotic cells [10, 11]. The eukaryotic dynamin-1 was the first dynamin protein studied, and it is involved in endocytosis by assembling into large helical structures around the neck of budding vesicles. This assembly is followed by GTPase activity, which releases the vesicles from the plasma membrane [12–15]. The other two isoforms, dynamin-2 and dynamin-3, which share 80% sequence similarity, mainly differ in their proline-rich domains and expression profiles [16]. Eukaryotic DLPs are classified as either membrane fission or membrane fusion DLPs based on their biological functions [17]. Mitochondrial division by Drp1 and peroxisomal division by DRP3A are examples of fission DLPs [18, 19], while membrane-fusing DLPs, such as Fzo1 mitofusins and Mgm1/OPA1, fuse the outer and inner mitochondrial membranes, respectively, and Atlastins fuse membranes of the endoplasmic reticulum [20–22]. A fascinating class of DLPs, called Mx proteins, have been shown to have a protective effect against a wide range of DNA and RNA viruses, making them part of the eukaryotic innate immune response [23]. However, their mode of action in anti-viral activity is still largely unclear [24].

Bacterial dynamin-like proteins (BDLPs), a subclass of DLPs, were first predicted in 1999, and since then, several hundred bacterial species have been identified to possess DLPs [25]. BDLPs gained attention when the full-length crystal structure of BDLP1 from *Nostoc punctiforme* was reported [26]. While the exact function of BDLP1 in *N. punctiforme* remained unclear, it demonstrated ordered self-assembly on liposome surfaces, reshaping them into highly curved tubules while disrupting the outer lipid layer [7]. Similarly, a SynDLP in the cyanobacterium *Synechocystis* showed similar self-assembly into high-molecular-mass oligomers on negatively charged membrane lipids [27]. BDLPs have been found in both Gram-positive and Gram-negative bacteria. Some BDLPs, such as DynA and DynB from *Streptomyces venezuelae*, a filamentous bacterium, have been shown to play a role in cell division during sporulation [28] Other BDLPs, such as IniA and IniC in *Mycobacterium tuberculosis* and LeoA in *Enterotoxigenic Escherichia coli*, are involved in extracellular vesicle release [29, 30], while the BDLP from *Campylobacter jejuni* is speculated to fuse biological membranes [31, 32].

DynA, a two-headed BDLP in *Bacillus subtilis*, was first described in 2011 and found to have two connected dynamin-like domains, D1 and D2, suggesting it resulted from gene duplication and fusion events [33, 34]. In vitro studies have revealed that DynA can tether and fuse membranes in trans. The D1 domain of DynA is crucial for membrane tethering and fusion, while the D2 domain facilitates the process [33, 35] GTP binding facilitates full membrane fusion and likely DynA disassembly, but is not required for membrane tethering and lipid mixing [33, 35]. Further research has shown that DynA is involved in a complex stress response pathway that protects the plasma membrane’s integrity. Deletion of *dynA* from *B. subtilis* renders the bacteria more sensitive to environmental stresses, such as pore-forming antibiotics and phages [36]. The protein exhibits high lateral mobility in the plasma membrane but forms large, confined clusters in response to the pore-forming antibiotic nisin, suggesting its recruitment to sites of membrane damage [36, 37]. This protective effect of DynA was further studied using the smallest *B. subtilis* phage, Ф29. DynA-deficient strains exhibited increased susceptibility to phage infection, as shown by larger plaque formations and extensive lysis in liquid cultures. Interestingly, DynA-overexpressing strains showed significantly less and delayed host cell lysis at the population level. During phage infection, DynA formed large immobile clusters in the later stages, indicating its role in blocking phage progeny release by preventing membrane rupture [37]. DynA is the first reported BDLP with a critical role in bacterial stress response and innate immunity.

This study focuses on characterizing the membrane-binding site within the D1 domain of DynA. Using a DynA AlphaFold model as a template, we identified a candidate membrane-binding site comprising key positively charged lysine residues (K360 and K367) and hydrophobic phenylalanine residues (F363, F364, and F365). These residues play a significant role in facilitating interaction with biological membranes. To investigate the functional relevance of this membrane-binding site, we engineered a series of point mutations in DynA. Our findings reveal that the mutants DynA F363A, F364A, and F365A, as well as DynA K360A and K367A, exhibit a complete loss of membrane-binding capability, as demonstrated by liposome sedimentation assays. In contrast, a mutant engineered with conservative substitutions, DynA F363W, F364W, and F365W, maintained its membrane-binding ability, suggesting that structural integrity around these residues is crucial for DynA’s membrane interaction. Furthermore, we employed electron microscopy to visualize DynA’s binding to lipid membranes. Wild-type DynA was observed to envelop liposomes, inducing liposome clustering indicative of successful membrane interaction, while the mutants displayed a stark absence of binding, further confirming our biochemical findings. The in vivo relevance of DynA’s membrane-binding capability was assessed through phage infection assays. We found that the membrane-binding mutants exhibited significantly increased susceptibility to lysis compared to wild-type strains, highlighting the importance of this interaction in conferring phage resistance. Moreover, our experiments demonstrated that the D1 domain alone could effectively mediate this protective function when expressed in *B. subtilis*, emphasizing its pivotal role.

## Material and Methods

**Table 1.** Bacterial strains used in this study.

| Strain | Genotype | Source/Reference |
| --- | --- | --- |
| ***Bacillus subtilis* strains** |  |  |
| *Bacillus subtilis* 168 | *trpC2* | Laboratory collection |
| *Bacillus subtilis* 25152 | *trpC2, metBI O, xin-1, SPβ-(erm)* | Laboratory collection |
| FBB002 | *trpC2, dynA::tet* | [33] |
| FBB018 | *trpC2, amyE::Pxyl-dynA-gfp spc* | [33] |
| SSB004 | *trpC2, amyE::Pxyl-dynA-F363A-F364A-F365A-gfp spc dynA::tet* | This study |
| SSB005 | *trpC2, amyE::Pxyl-dynA-K360A-K367A-gfp spc dynA::tet* | This study |
| LJ-B05 | *trpC2, amyE::Pxyl-D1-yfp-spec dynA::tet* | This study |
| LJ-B06 | *trpC2, amyE::Pxyl-D2-gfp-spec dynA::tet* | This study |
| SSB006 | *trpC2, amyE::Pxyl-D1-F363A-F364A-F365A - yfp-spec dynA::tet* | This study |
| SSB007 | *trpC2, amyE::Pxyl-dynA-F363W-F364W-F365W-gfp spc dynA::tet* | This study |
| ***Escherichia coli* strains** |  |  |
| BL21 (DE3) | $F^{-} omp^T hsd S_B (r_B^{-}, m_B^{-}) gal dcm$ (DE3) | Novagen |
| SSE001 | BL21 (DE3) pET16b_DynA-His6 | This study |
| SSE002 | BL21 (DE3) pET16b_DynA<br>F363A, F364A, F365A -His6 | This study |
| SSE003 | BL21 (DE3) pET16b_DynA<br>K360A, K367A -His6 | This study |
| SSE004 | BL21 (DE3) pET16b_DynA<br>F363W, F364W, F365W -His6 | This study |
| SSE005 | BL21 (DE3) pET16b_DynA<br>K56A, K625A -His6 | This study |

### Growth and isolation of phages

Ф29 phages were obtained from the German Collection of Microorganisms (DSMZ GmbH). The host strain *B. subtilis* 25152 was cultured to mid-log phase in LB medium at 37 °C with continuous shaking. High-titer phages (approximately 10^8–10^10 PFU) were then added to the bacterial culture. The mixture was incubated at 37 °C for 6 hours with shaking. After incubation, the phage-host mixture was centrifuged at 3,800 x g for 20 minutes to separate cell debris. The phage supernatant was filtered through 0.2 μm pore filters and stored at 4 °C.

### Phage purification by NaCl-PEG precipitation

One liter of the bacteria-phage mixture was centrifuged at 3,800 x g for 20 minutes to separate the phages from the cell debris. The supernatant was then filtered, and solid NaCl was gradually added to a final concentration of 1 M, stirring until fully dissolved. Next, 1% solid PEG 8000 (Millipore Sigma, Burlington, MA, USA) was slowly added to the supernatant, and the mixture was stirred continuously for 30 minutes. The phage mixture was pelleted by centrifugation at 15000 × g for 10 minutes at 4 °C. The supernatants were gently decanted without disturbing the pellet. The pellets were then completely resuspended in 20 mL of SM Buffer (100 mM NaCl, 8 mM MgSO₄·7H₂O, 50 mM Tris-Cl, pH 7.5). The phages in SM buffer were centrifuged again at 15000 × g for 10 minutes at 4 °C. The purified phages were separated from the pellets and stored at 4 °C.

### Phage infection assay in liquid medium

Overnight culture of *B. subtilis* and strains that expressed DynA mutant proteins were diluted to an OD_600_ of 0.1 and grown until optical density reached approximately 0.3. The cultures were then infected with Φ29 phage at a multiplicity of infection (MOI) of 1, and optical density was measured every 10 minutes using Infinite200 PRO (Tecan, Grödig, Austria) over a period of 14 hours at 37°C with continuous shaking.

### Quantitative plaque assay

Overnight cultures of *B. subtilis* were diluted to an OD_600_ of 0.1 in fresh LB medium and grown until they reached an OD_600_ of 0.5–1.0. The purified phages were serially diluted tenfold (from 10^1 to 10^10) in gelatin-free SM buffer. For each dilution, 100 µL of the phage solution was mixed with 1 mL of bacterial culture and incubated at room temperature for 10 minutes to facilitate phage attachment. Each bacterial-phage mixture was then combined with 4 mL of overlay agar (containing 0.4% agar in LB medium) and poured onto LB agar underlay plates (1% agar). The plates were incubated at 24 °C for 18 hours, after which plaques were observed.

### Protein purification

Heterologous expression of DynA-His₆ and its mutants was conducted in *E. coli* BL21 (DE3) strains in LB medium supplemented with the appropriate antibiotics (carbenicillin at 100 μg/mL). Protein expression was induced by adding IPTG to a final concentration of 0.5 mM when the OD_600_ reached approximately 0.7. The cells were incubated overnight at 18 °C. Cultures were harvested by centrifugation at 6500 x g for 15 minutes at 4 °C, washed with T2 buffer (50 mM Tris, 200 mM NaCl, 20 mM imidazole, 10% glycerol, pH 8.0), and then centrifuged again at the same speed and temperature. The cell pellets were frozen and stored at -80 °C.

The frozen pellets were resuspended in cold T2 buffer and supplemented with lyophilized DNase I and Protease Inhibitor Cocktail (cOmplete™, EDTA-free, Roche 04693132001), along with 0.7% Triton X-100. The suspension was passed through a French Pressure Cell (Amico) to disrupt the cells, with the process repeated three times at an inner pressure of 20,000 psi. The resulting suspension was centrifuged at 15,000 × g at 4 °C using a Beckman Coulter Avanti J-25 centrifuge with a JA-10 rotor and Falcon adapters to remove cell debris. The supernatant was transferred to a fresh Falcon tube and mixed with T2-buffered Ni-NTA agarose (Qiagen), followed by incubation at 4 °C overnight with continuous mixing.

The following day, the bound protein was extensively washed with T2 buffer and eluted using T5 buffer (50 mM Tris, 500 mM NaCl, 1 M imidazole, 10% glycerol, pH 8.0) for one hour at 4 °C with continuous shaking. The proteins were concentrated as needed using Amicon filter devices with the appropriate molecular weight cut-off (Millipore; Merck). The eluted protein was reduced with 1 mM DTT and subjected to size-exclusion chromatography using a Superose 6 Increase column (Cytiva) in 50 mM Tris, 500 mM NaCl, 10% glycerol, pH 8.0. Finally, the protein concentration was determined using the Bradford assay and stored at -80 °C until further use.

### GTP hydrolysis assays

GTP hydrolysis was analyzed in continuous reactions using the EnzChek Phosphate Assay Kit (E-6646, Molecular Probes Inc.), following the manufacturer’s instructions. Reaction volumes were reduced to 100 μL and assayed in flat-bottom 96-well plates (Greiner-UV-Star 96-well plates). Reactions were measured at 37 °C, continuously, every minute, over a 3-hour time course in a Tecan Infinite 200 Pro with the Tecan i-control v.2.0 software. Each reaction contained 5 μM DynA or respective DynA mutants, buffered in 50 mM Tris, 200 mM NaCl, pH 8.0. Before starting the reactions with 2 mM Mg-GTP, the reactions were pre-incubated for 10 minutes to eliminate phosphate contamination. Data were analyzed with Excel (normalized, with subtraction of GTP auto-hydrolysis and subtraction of the no-substrate control, and linear regression was used to determine the hydrolysis rate per minute). Data visualization and further analysis were performed with GraphPad Prism version 5.03 for Windows (GraphPad Software).

### Liposome sedimentation assay

A thin film of Avanti total *Escherichia coli* polar lipids was dissolved in chloroform and dried under a nitrogen stream for 3 hours. The dried film was then rehydrated in 50 mM Tris, 10% glycerol, pH 7.1, at 37 °C for one hour. The resulting suspension was diluted to 2 mg/mL in assay buffer (50 mM Tris, 200 mM NaCl, 10% glycerol, pH 7.4) and extruded 40 times through a 400 nm pore size filter using a mini extruder. Two micromolar DynA protein was incubated with 1 mg/mL liposome solution at 25 °C. The mixture was then ultracentrifuged at 90,000 × g for 20 minutes. Nucleotides were used at a 1 mM concentration with 5 mM MgSO₄. The supernatants were carefully separated from the pellets, and protein presence was analyzed using SDS-PAGE.

### Liposome preparation for electron microscopy

A thin film of Avanti total *Escherichia coli* polar lipids was dissolved in chloroform and dried under a nitrogen stream for at least 1 hour. The lipids were further dehydrated in a vacuum dryer for another 4 hours. The lipids were then rehydrated in 50 mM Tris, 10% glycerol, pH 7.1, at 37 °C for 3 hours. The resulting suspension was diluted to 2 mg/mL in assay buffer (50 mM Tris, 200 mM NaCl, 10% glycerol, pH 7.4) and extruded 40 times through a 200 nm pore size filter using a mini extruder.

### Electron microscopy

For the liposome membrane-binding assay, 25 µM DynA/mutant protein was incubated with 1 mg/mL liposomes in the absence of the GTP nucleotide at 24 °C for 10 minutes. The samples were immediately analysed by negative staining transmission electron microscopy. To this end, copper, 400 square mesh, formvar-coated TEM grids were glow discharged for 60 s at 0.6 mbar air pressure and 10 mA glow current using a Safematic CCU-010 unit, and then incubated with 5 μl of the liposome suspension for 4 min. Grids were washed shortly on 7 drops of water and embedded in a thin layer of 1.8% methyl cellulose with 0.2% uranyl acetate. Samples were imaged with a Tecnai G2 Spirit BioTwin transmission electron microscope (FEI / now Thermo Fisher Scientific) operated at 80 kV using TEM User interface (v. 4.2) and equipped with a LaB6 filament. Images were recorded with a MegaView III G2 CCD camera, using iTEM v.5 software (both Olympus Soft Imaging solutions / now EMSIS).

### Statistical analysis

We used the SPSS software (SPSS Inc., Chicago, Ill., USA) to calculate statistical significances

## Results

### Membrane-binding site of DynA

DynA consists of two structurally similar subdomains, D1 and D2, both containing GTP-binding motifs essential for GTP hydrolysis [33]. However, previous studies indicated that the D1 domain is primarily responsible for membrane tethering and fusion, as demonstrated by in vitro assays [33, 35]. In our initial screening of potential membrane-binding regions within the D1 domain, we identified a candidate site using an AlphaFold v2.0 generated model [38]. The model shows two globular GTPase domains located in close proximity to each other. Each of these GTPase domains are connected to an alpha helical stalk region. The stalk of D1 domain exhibits an extended conformation while the stalk of D2 domain adopts a more folded confirmation forming an overall compact structure of DynA. The extended tip of the D1 stalk has two positively charged lysine residues at positions 360 and 367, along with three consecutive hydrophobic phenylalanine residues at positions 363, 364, and 365, making this region electrostatically and hydrophobically ideal to interact with lipid membranes (Figure 1). Moreover, this specific arrangement of residues is only present within the D1 stalk while being absent in the D2 domain.

**Figure 1:**
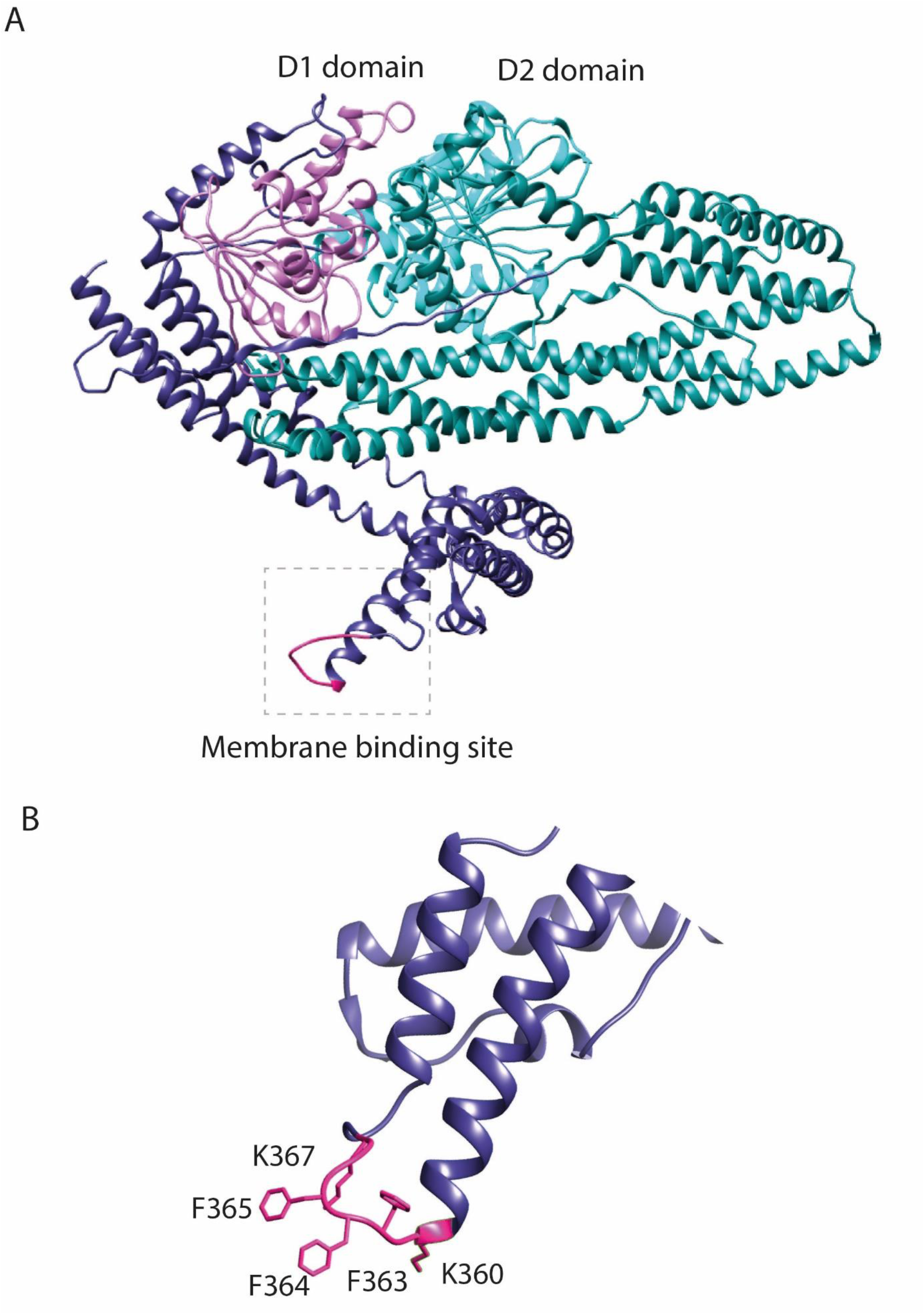
AlphaFold v2.0 model of *B. subtilis* bacterial dynamin like protein DynA. (A) DynA possesses two dynamin domains D1 (purple) and D2 (cyan) with their GTP-binding domains. The D1 domain possesses a putative membrane-binding site (pink) at the end of the extended stalk. (B) A close-up view of the putative membrane-binding site highlighting the arrangement of key lysine residues at position 360 and 367 and three phenylalanine residues at position 363,364 and 365.

### DynA F363A, F364A, F365A and DynA K360A, F367A variants lack membrane-binding

To further examine the region between residues K360 and K367 as a potential membrane-binding site, we introduced three-point mutations to create the DynA F363A, F364A, F365A mutant protein, and two-point mutations to generate the DynA K360A, K367A mutant protein. Additionally, we engineered a third mutant, DynA F363W, F364W, F365W, with conservative substitutions, to assess whether the membrane-binding capability of DynA is retained under these conditions. We also included the previously described DynA K56A, K625A mutant, which lacks GTPase activity, as a control in our experiments [33].

The DynA protein and its four mutants were heterologously expressed with a poly-His tag in *E. coli* and purified in a two-step chromatography (see Materials and Methods section). All proteins were purified to homogeneity and subsequently used for in vitro assays. DynA and all its mutants exhibited basal GTPase activity that followed Michaelis–Menten kinetics (Figure 2). We interpreted the presence of GTPase activity as an indication that the generated mutant variants of DynA contained their overall fold.

**Figure 2:**
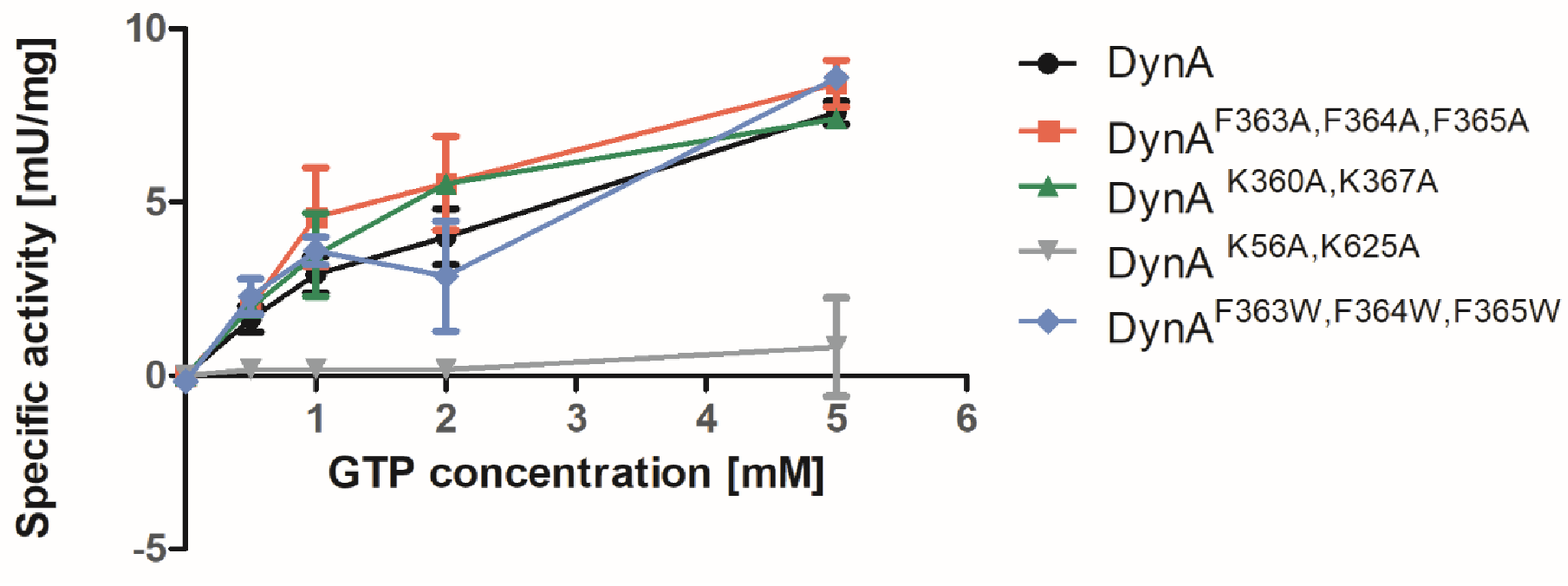
GTPase activity of 5 μM DynA and DynA mutants. Error bars indicate the standard deviation of three independent experiments.

Next, we evaluated the membrane-binding ability of DynA and its mutant proteins using liposome sedimentation assays. Small, unilamellar vesicles, approximately 400 nm in diameter, were prepared and incubated at 24°C with the protein of interest. The protein-liposome mixture was then subjected to ultracentrifugation, and the resulting pellets and supernatants were separated and analyzed for protein presence using SDS-PAGE (Figure 3A).

**Figure 3:**
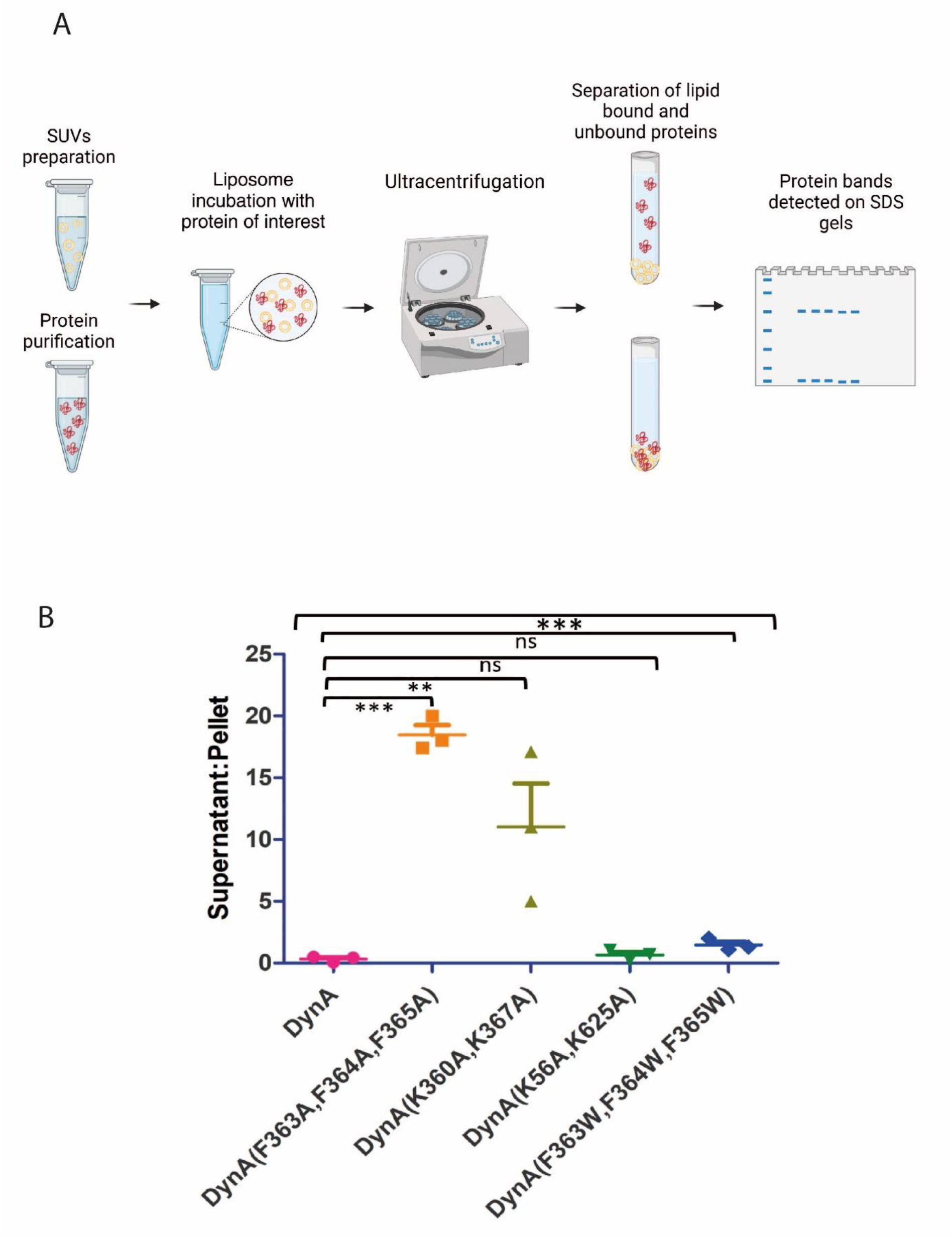
Liposome sedimentation assays of WT DynA and DynA mutants. (A) Graphical illustration of the liposome sedimentation assay (B) Supernatant to pellet ratios of DynA, DynA F363A, F364A, F365A, DynA K360A, K367A, DynA K56A, K625A and DynA F363W, F364W, F365W variants, respectively. WT DynA exhibits a low supernatant to pellet ratio indicating effective membrane-binding. While both mutant proteins DynA F363A, F364A, F365A and DynA K360A, K367A remained in the supernatants, the conservative mutant DynA F363W, F364W, F365W retained the membrane-binding. Statistical analysis is based on One-way Anova and Post-hoc Tukey’s Honestly Significant Difference (HSD) test (*, P value less than 0.05; **, P value less than 0.01; ***, P value less than 0.001). Error bars depict standard deviation of three independent experiments.

The results from the liposome sedimentation assay demonstrated a complete loss of membrane-binding ability in both the DynA F363A, F364A, F365A and DynA K360A, K367A mutants compared to the wild-type DynA. In contrast, the GTPase mutant DynA K56A, K625A, while lacking GTPase activity, retained its membrane-binding capability, as previously described [33]. Moreover, the mutant with conservative substitutions, DynA F363W, F364W, F365W, was able to bind liposomes (Figure 3B). These data provide compelling evidence that the region between residues K360 and K367 constitute the membrane-binding domain (paddle domain) of DynA.

### Electron microscopy reveals loss of membrane binding in DynA F363A, F364A, F365A and DynA K360A, K367A

To gain further information on the interaction of DynA with liposome membranes, we employed transmission electron microscopy to gain visual insights into the DynA membrane interaction. When 200 nm liposomes were incubated with 25 µM DynA, the protein was observed to completely envelop the liposomes, and several liposomes aggregated to form clusters held together by DynA. Arrowheads indicate the electron-dense layer representing the contrasted DynA protein on the surface of the cup-shaped liposomes. On the contrary, the DynA F363A, F364A, F365A mutant was unable to attach to or aggregate any liposome vesicles. Similarly, the DynA K360A, K367A mutant showed no binding affinity for the liposomes; however, liposome destabilization and degradation were evident in the presence of this protein (Figure 4).

**Figure 4.**
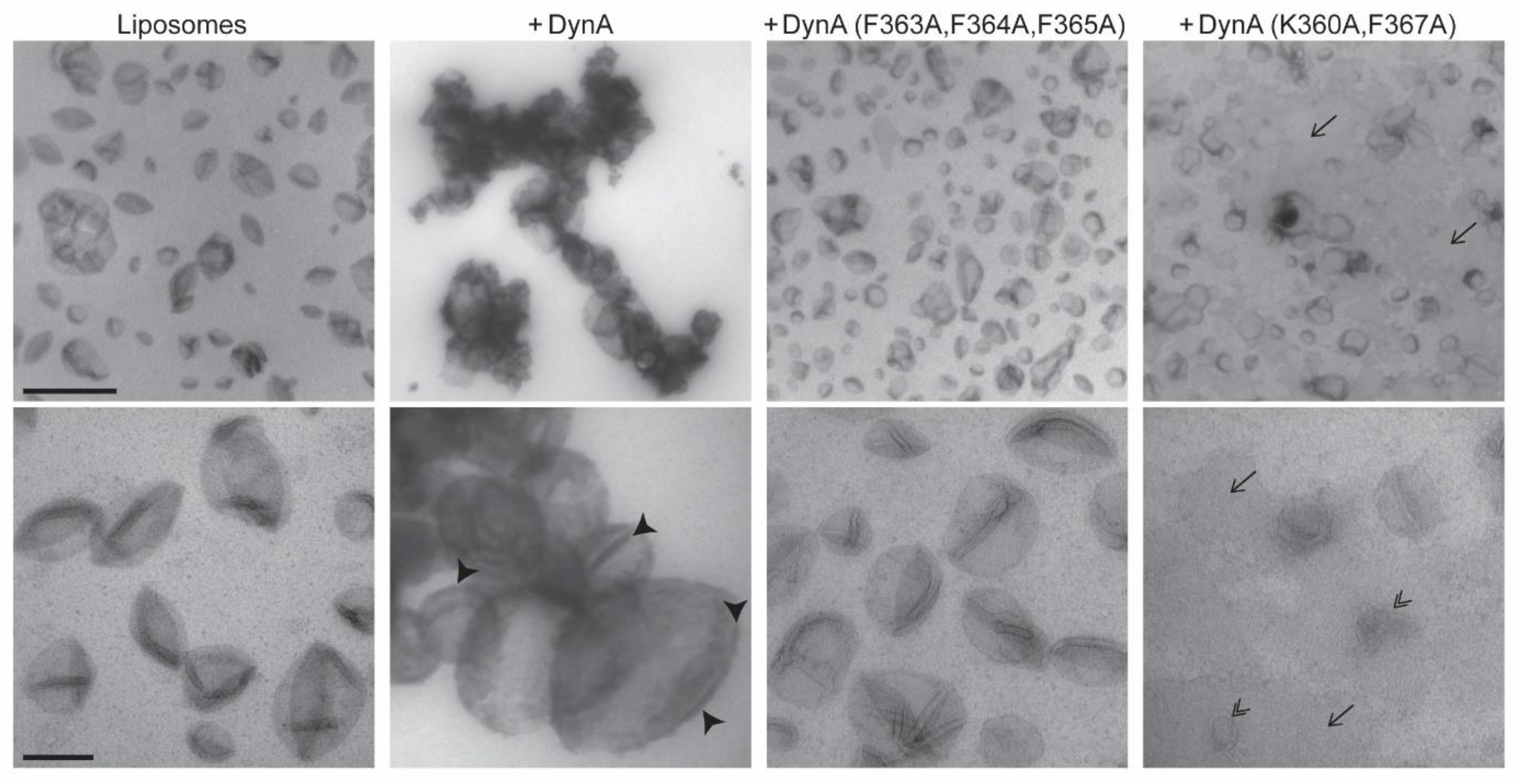
Liposomes incubated with 25 µM DynA in the absence of the nucleotide were examined by transmission electron microscopy. Wild type DynA binds the liposome membrane, leading to tethering and clumping of liposomes. Arrowheads point to the electron dense layer representing contrasted DynA protein on the surface of cup-shaped liposomes. In contrast, in the presence of the DynA F363A, F364A, F365A mutant, liposomes remain dispersed. In the presence of the DynA K360A and K367A mutant, liposomes were destabilized (double arrowheads) and insoluble lipids accumulated on the film (arrows). Liposomes were embedded in 0.2% visualized by TEM after embedding in 1.8% methyl cellulose with 0.2% uranyl acetate. Scale bar: 1 µm (upper panel), 200 nm (lower panel).

### DynA’s membrane-binding is essential for phage protection

Phage Ф29 produces small plaques on a lawn of *B. subtilis* cells [39]. Previous studies from our laboratory have demonstrated a unique protective effect of DynA during Ф29 phage infection. Cells lacking DynA exhibited significant host cell lysis in presence of phages [36], as shown by plaque assays and infection assays in liquid cultures, suggesting a protective effect of DynA. This protective effect of DynA becomes more pronounced with DynA overexpression, resulting in a substantial reduction in host cell lysis [37]. Our objective was to determine whether the membrane-binding capability of DynA is essential for its in vivo protective effect against phage infection. To investigate this, strains were generated that expressed various DynA mutant proteins as a sole DynA copy ectopically from the *amyE* locus under a xylose-inducible promoter. In vivo assays in liquid medium and on agar plates were performed to test the resistance of these strains to Ф29 phage lysis.

First, we analyzed the lysis behavior of a strain expressing the mutant DynA F363A, F364A, F365A (strain SSB004). The DynA mutant was expressed as a GFP fusion to allow microscopic inspection. Exponentially growing cultures of wild-type *B. subtilis*, the DynA deletion strain (*B. subtilis*, Δ*dynA*), the DynA membrane mutant strain (DynA F363A, F364A, F365A), and the DynA overexpression strain (DynA++) were infected with a multiplicity of infection (MOI) of 1, and optical density was measured continuously in a plate reader over a period of 14 hours. The strains expressing DynA F363A, F364A, F365A membrane-binding mutant as sole copy exhibited significantly increased cell lysis, similar to the deletion strain, indicating that the absence of DynA membrane-binding renders cells more susceptible to lysis upon phage infection (Figure 5Ai). A similar effect was observed in plaque assays, where the lawn of the membrane-binding mutant strain (DynA F363A, F364A, F365A) infected with Ф29 displayed much larger plaques and twice the number of plaques compared to wild-type *B. subtilis* 168 (Figure 5Aii).

**Figure 5:**
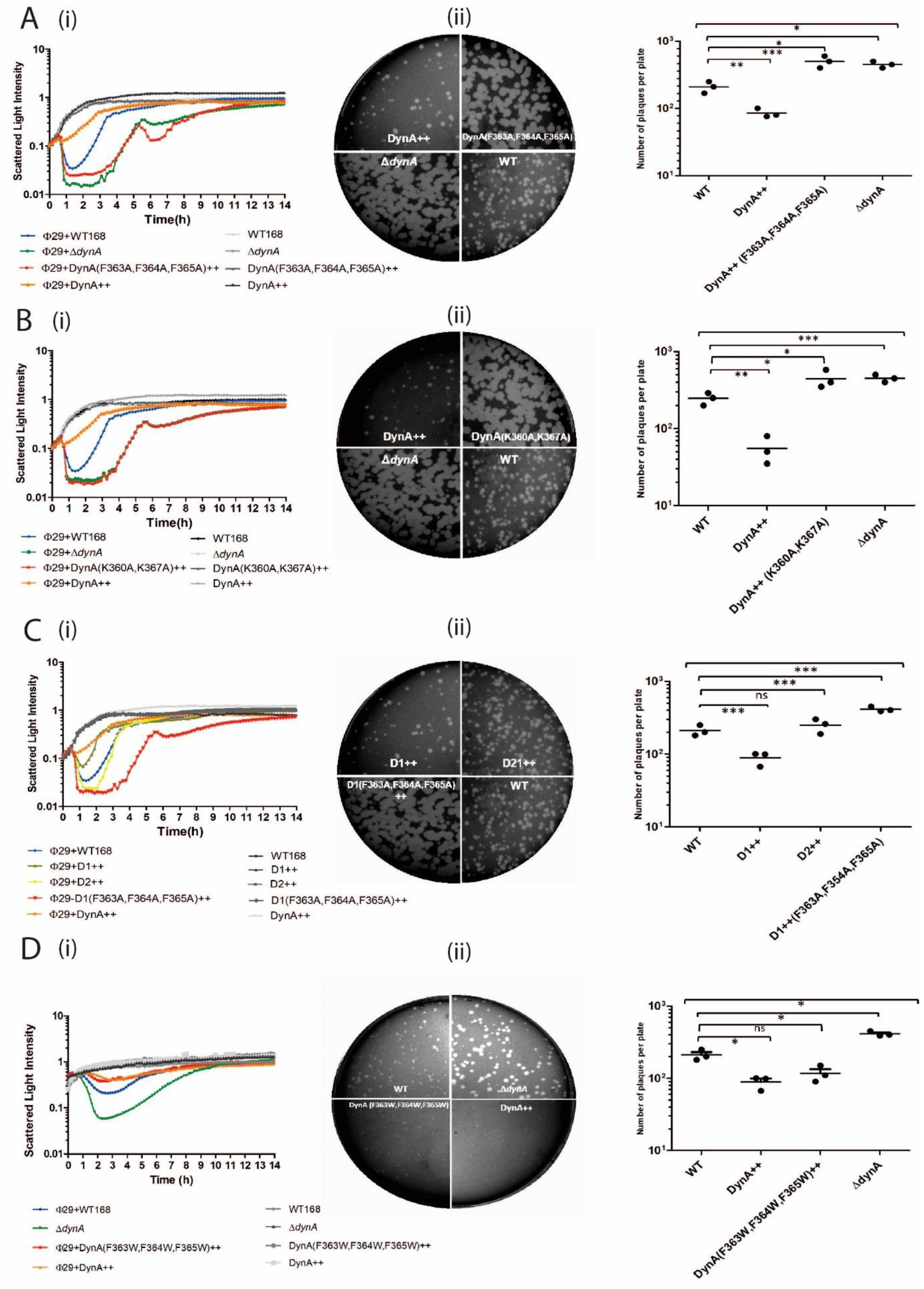
Membrane-binding of DynA is essential for bacterial innate immunity. The lytic effect of Ф29 infection (MOI=1) on different *B. subtilis* strains was tested (i) by measuring the optical densities of liquid growth cultures in a plate reader, and (ii) by ɸ29 plaque formation: photos of agar plates are displayed in the middle column; quantitative analysis is presented in the right column. Statistical analysis is based on One-way Anova and Post-hoc Tukey’s Honestly Significant Difference (HSD) test (*, P value less than 0.05; **, P value less than 0.01; ***, P value less than 0.001). Error bars indicate the standard deviation of three independent experiments. The following bacterial strains were used for comparison: wild type *B. subtilis* 168 (WT), DynA-deficient strain (Δ*dynA*), DynA overexpressing strain (DynA++). (A) The DynA F363A, F364A, F365A ++ strain shows a similar pattern of cell lysis as the DynA deficient strain (i), and larger plaques compared to WT. (B) The DynA K360A, K367A ++ mutant strain shows a similar pattern of cell lysis as that of DynA deficient strain (i), and larger plaques compared to WT (ii). (C) Cells expressing only the D1 domain of DynA show resistance to the phage infection similar to the overexpressing strain, while the D2 domain alone is ineffective in preventing host cell lysis. Mutation in the D1 F363A, F363A, F365A domain renders the community sensitive to phage infection (i, ii). (D) DynA conservative mutant shows protective effect against ɸ29 infection (i,ii).

Next, we analyzed the second membrane-binding mutant strain, DynA K360A, K367A (strain SSB005), again expressed as a GFP fusion. The infection trend for this mutant mirrored that of the phenylalanine triple mutant described before, showing a higher degree of cell death when exponentially growing cells were infected with Ф29 at a MOI of 1 compared to the wild-type strain. In contrast, the DynA overexpression strain (DynA++) demonstrated resistance to cell lysis, consistent with prior observations (Figure 5Bi). The plaque assay for the membrane mutant (DynA K360A, K367A) strain revealed larger and more numerous plaques, similar to those seen in the DynA deletion strain, underscoring the importance of positively charged residues for membrane-binding (Figure 5Bii).

Previous work based on liposome sedimentation assays has shown that the D1 domain of DynA exhibits in vitro membrane-binding, while the D2 domain does not possess any binding ability [33]. We aimed to determine if the D1 domain alone could fulfill the functional role of DynA in vivo. To this end, we constructed a *B. subtilis* strain expressing only the D1 domain of DynA, fused to YFP (strain: LJ-B05). We compared the D1-expressing strain to a D2-expressing strain (strain: LJ-B06) by simultaneously infecting both with Ф29 phages during their exponential growth phase at a MOI of 1. The D1-expressing strain showed resistance to cell lysis, while the D2-expressing strain exhibited sensitivity, with lysis behavior similar to that of the wild-type strain. Additionally, we created a DynA D1 F363A, F364A, F365A mutant (SSB006), again expressed as a GFP fusion. This D1 membrane-binding mutant was sensitive to cell lysis upon infection with Ф29 phages (Figure 5Ci). The plaque assay results were consistent with the infection assays in liquid cultures, further confirming that membrane-binding is integral to the D1 domain of DynA and that this binding is essential for its protective effect against phages (Figure 5Cii).

A DynA F363W, F364W, F365W conservative mutant strain (strain SSB007) with hydrophobic side chains was developed to assess whether we can conserve the membrane-binding of DynA in vivo. This mutated DynA variant was again expressed as a GFP fusion. Exponentially growing cultures of wild-type *B. subtilis*, DynA deletion strain (*B. subtilis*, Δ*dynA*), DynA conservative mutant strain (DynA F363W, F364W, F365W), and DynA overexpression strain (DynA++) were infected with Ф29 phages at a MOI of 1, and optical density was measured. The DynA F363W, F364W, F365W conservative mutant showed significantly less cell lysis, similar to the DynA overexpression strain (DynA++), further confirming that phenylalanine residues at positions 363, 364, and 365 are integral parts of the membrane-binding site of DynA (Figure 5Di). Similar results were obtained by plaque assays (Figure 5Dii). In summary these data confirm that membrane-binding of the D1 part of DynA plays a crucial role in phage defense.

## Discussion

The bacterial dynamin-like protein DynA in *B. subtilis* is part of a novel phage protection system that acts as a critical last line of defense against phage infection. By delaying host cell lysis and preventing the immediate release of phage progeny after replication and assembly, DynA helps to mitigate the rapid spread of phages through the bacterial population [37]. To fully understand the molecular mechanisms underlying this protective effect, it is essential to explore the complex structure of DynA and the roles of its individual domains and domains. DynA is a membrane-associated protein with an approximate molecular weight of 137 kDa. It consists of two domains, D1 and D2, and the holoenzyme dimerizes in vitro. Both domain contain independent GTP-binding domains. Each GTP-binding domain features a central P-loop motif, which includes an essential lysine residue crucial for nucleotide binding. Mutation of both P-loops (K65A and K625A) in DynA abolishes GTP binding and hydrolysis, as shown in previous studies [33]. Notably, sequence analysis of DynA reveals the absence of traditional transmembrane domains [33]. However, earlier studies also revealed that DynA and DynA D1 co-sediment with liposomes, while DynA D2 does not exhibit membrane-binding. This suggests that membrane tethering is governed exclusively by the D1 domain, whereas the D2 domain plays a crucial role in stabilizing the DynA complex [33, 35]. Interestingly, DynA and a distantly related antiviral dynamin Mx1 share a unique functional similarity. Neither the DynA membrane tethering nor the Mx1 interaction with the viral ribonucleoprotein requires GTP hydrolysis. [33, 40]. However, both proteins require GTP hydrolysis for their antiviral protective effect [37, 41].

In our study, we used an AlphaFold model to investigate the potential membrane-binding site within the D1 domain of DynA. This analysis identified two positively charged lysine residues at positions 360 and 367, alongside three consecutive hydrophobic phenylalanine residues at positions 363, 364, and 365 in the extended stalk region of the D1 domain as a potential membrane-binding site. Previous studies on human MxA proteins have documented the importance of lysine residues at the tip of the stalk region in facilitating electrostatic interactions with cellular membranes [42, 43].

To test the potential membrane-binding role of this region in DynA, we introduced mutations in the D1 domain. Specifically, we generated the DynA F363A, F364A, F365A mutant and the DynA K360A, K367A mutant. Both mutants demonstrated a complete loss of membrane-binding in liposome sedimentation assays. Interestingly, the DynA F363W, F364W, F365W mutant, which incorporated conservative substitutions, retained the ability to co-sediment with liposomes. These findings suggest that the arrangement of positively charged lysine residues surrounding hydrophobic phenylalanine residues in the D1 stalk region is essential for DynA’s membrane-binding capability. Similarly, hydrophobic residues in the paddle domain of BDLP from *N. punctiforme* have been shown to be involved in membrane-binding, where mutations in phenylalanine and leucine residues disrupted lipid association [7]. Therefore, the paddle domain of BDLP and DynA share structural and functional similarities.

Next, we aimed to investigate the importance of DynA’s membrane-binding in vivo. Previous studies have demonstrated that DynA plays a unique protective role against phage infection by delaying host cell lysis [37]. However, the molecular mechanism of DynA action in bacterial innate immunity remained unclear. To determine if this protective effect is mediated through direct interactions with the plasma membrane, we compared the lysis behavior of *B. subtilis* strains expressing either the DynA F363A, F364A, F365A or DynA K360A, K367A mutant, as the sole DynA copy, to that of the wild-type strain during phage infection. Both membrane-binding mutants exhibited significantly increased cell lysis during Ф29 phage infection compared to wild-type cells, confirming that DynA’s protective effect against phage infection is lost when its membrane-binding ability is compromised. These results suggest that DynA directly interacts with the membrane, forming a physical barrier to prevent rupture during phage induced host cell lysis. Furthermore, a strain expressing only the D1 domain of DynA exhibited reasonable protection against phage infection, while a strain expressing the D1 F363A, F364A, F365A mutant failed to protect against phage infection, further reinforcing the critical role of membrane-binding in DynA-mediated phage protection.

DynA protective phenotype against phages is similar to Mx dynamins that are a part innate immune response in eukaryotes by possessing antiviral activity. Both dynamins share remarkable structural and functional similarities.[36, 43]. However, unlike MxA, DynA shows no direct interaction with the phage particle so far, but the possibility persists. Two interesting bacterial dynamins LeoA of enterotoxigenic *E. coli* (ETEC) and IniA from *Mycobacterium tuberculosis* have been implicated in membrane fission and release of membrane vesicles [44, 45]. IniA has been speculated to assemble on the periplasmic side of the plasma membrane to bud membrane vesicles encapsulating anti-tuberculosis drugs thus contributing to resistance against anti-tuberculosis drugs [46]. In contrast DynA binds the membrane from the cytoplasmic site and hence production of vesicles that could act as phage decoys is unlikely. However, there is a possibility that DynA protects against phages acts by forming large assemblies on the plasma membrane where it captures and releases phage progeny that remain stuck at the lysing cell debris, thus preventing disruptive bursts and phage progeny dispersal.

In summary, this study provides a detailed understanding of the structural and functional characteristics of DynA, linking its membrane-binding ability to its protective role against phage infection. By elucidating the specific interactions at the membrane-binding site within the D1 domain of DynA, we provide the first evidence that DynÁs function in bacterial innate immunity requires its membrane-binding activity.

## Acknowledgements

We wish to acknowledge Manuela Weiss, Ekaterina Karnaukhova and Sara Baur for their assistance in this study, Lijun Guo for help with strain construction, and all members of the Bramkamp laboratory for critical discussions. The authors acknowledge co-funding from the Higher Education Commission of Pakistan (HEC) “Overseas Scholarship Scheme” (57558560) and DAAD (German Academic Exchange Service). We also acknowledge support by funding from the Deutsche Forschungsgemeinschaft (BR2915/13-1 to M.B.).

